# Purinergic P2Y2 Receptor-Induced Activation of Endothelial TRPV4 Channels Mediates Lung Ischemia-Reperfusion Injury

**DOI:** 10.1101/2023.05.29.542520

**Authors:** Maniselvan Kuppusamy, Huy Q. Ta, Hannah N. Davenport, Abhishek Bazaz, Astha Kulshrestha, Zdravka Daneva, Yen-Lin Chen, Philip W. Carrott, Victor E. Laubach, Swapnil K. Sonkusare

**Author notes:** Corresponding Author: Swapnil K. Sonkusare, Ph.D. 409 Lane Rd; room 6051A Charlottesville, VA 22901.

## Abstract

Lung ischemia-reperfusion injury (IRI), characterized by inflammation, vascular permeability, and lung edema, is the major cause of primary graft dysfunction after lung transplantation. We recently reported that endothelial cell (EC) TRPV4 channels play a central role in lung edema and dysfunction after IR. However, the cellular mechanisms for lung IR-induced activation of endothelial TRPV4 channels are unknown. In a left-lung hilar ligation model of IRI in mice, we found that lung IR increases the efflux of extracellular ATP (eATP) through pannexin 1 (Panx1) channels at the EC membrane. Elevated eATP activated elementary Ca^2+^ influx signals through endothelial TRPV4 channels through purinergic P2Y2 receptor (P2Y2R) signaling. P2Y2R-dependent activation of TRPV4 channels was also observed in human and mouse pulmonary microvascular endothelium in *ex vivo* and *in vitro* surrogate models of lung IR. Endothelium-specific deletion of P2Y2R, TRPV4, and Panx1 in mice had substantial protective effects against lung IR-induced activation of endothelial TRPV4 channels, lung edema, inflammation, and dysfunction. These results identify endothelial P2Y2R as a novel mediator of lung edema, inflammation, and dysfunction after IR, and show that disruption of endothelial Panx1–P2Y2R–TRPV4 signaling pathway could represent a promising therapeutic strategy for preventing lung IRI after transplantation.

## Introduction

The success of lung transplantation is limited by a high rate of primary graft dysfunction (PGD) due to ischemia-reperfusion injury (IRI) (*1, 2*). PGD is the leading cause of death in the first 30 days following lung transplant and is also an independent risk factor for chronic allograft dysfunction (*3, 4*). Lung IRI is a sterile inflammatory process characterized by robust oxidative stress, endothelial dysfunction, vascular permeability, edema, and alveolar damage after transplant (*5, 6*). While a number of signaling pathways are involved in lung IRI, it is clear that the cellular release of adenosine triphosphate (ATP) plays a prominent role (*1, 5*). Extracellular ATP (eATP) is an endogenous activator of purinergic receptor signaling. Therefore, targeting purinergic receptor-dependent mechanisms may be of therapeutic value in lung IRI. In the present study, we focused on the signaling mechanisms through which eATP and purinergic receptor signaling regulate vascular inflammation, edema, and dysfunction—the hallmarks of lung IRI.

Endothelial dysfunction and increased endothelial cell (EC) permeability are critical contributors to immune cell infiltration into pulmonary tissue and alveolar space, leading to lung edema after IR. EC pannexin 1 (Panx1) is an important source of eATP in the intact pulmonary endothelium under normal conditions (*7, 8*). We previously showed that endothelium-specific Panx1-knockout (Panx1_EC_^-/-^) mice are protected against vascular permeability, inflammation, and edema after lung IR (*9*), suggesting an important role for endothelial Panx1 and release of eATP in the pathogenesis of lung IRI. However, eATP-induced downstream signaling mechanisms that mediate lung IRI remain unknown. We recently showed that eATP, effluxed via endothelial Panx1 channels, activates endothelial P2Y2 purinergic receptors (P2Y2Rs) to regulate pulmonary arterial pressure under normal conditions (*8*). Therefore, we postulated that eATP-induced activation of endothelial P2Y2Rs mediates lung edema and dysfunction following lung IR.

Transient receptor potential vanilloid 4 (TRPV4) channels, which constitute a major Ca^2+^-entry pathway in pulmonary endothelium (*8, 10, 11*), have emerged as central players in the pathogenesis of different forms of lung injury (*12–15*). We provided the first evidence that endothelial TRPV4-knockout (TRPV4_EC_^-/-^) mice are protected against increased vascular permeability, lung inflammation, and edema after lung IR (*16*), suggesting that endothelial TRPV4 channels are critical mediators of lung IRI. However, whether increased activity of pulmonary endothelial TRPV4 channels contributes to lung IRI is not known. Recent studies provide evidence that, under healthy conditions, eATP activates endothelial TRPV4 channels through P2Y2R signaling (*8*). Therefore, we hypothesized that eATP, effluxed via endothelial Panx1 channels, activates endothelial P2Y2R–TRPV4 channel signaling to mediate lung inflammation, edema, and dysfunction in lung IRI.

To test our hypothesis, we investigated the role of endothelial Panx1–P2Y2R– TRPV4 signaling in lung IRI using tamoxifen-inducible, endothelium-specific P2Y2R,(*17*) Panx1 (*18*), and TRPV4 (*19*) knockout mice in combination with *in vivo*, *ex vivo*, and *in vitro* models of lung IRI. Our data show that eATP, derived by efflux via endothelial Panx1 channels, activates endothelial TRPV4 channels through P2Y2R signaling to promote lung inflammation and edema, and impair lung function in IRI. These data identify P2Y2R as a novel intermediate in an endothelial Panx1–P2Y2R–TRPV4 signaling axis and provide proof of principle that disrupting this pathway could be a potential treatment strategy for preventing IRI and PGD after lung transplantation.

## Results

### Lung IR increases the activity of TRPV4_EC_ channels via eATP

Based on our findings that TRPV4_EC_^-/-^ mice are protected against lung IRI,(*16*) we hypothesized that increased Ca^2+^ influx through TRPV4_EC_ channels mediates lung IRI. To test this, we used a well-established left-lung hilar-ligation model of IRI in mice (Fig. 1A) involving 1 hour of ligation (ischemia) followed by 2 hours of reperfusion (*9, 20*). After IR, mice display significant lung dysfunction, neutrophil infiltration, and increased levels of inflammatory cytokines in bronchoalveolar lavage (BAL) fluid (*16*). Control mice underwent sham surgery with no hilar ligation. The activity of individual Ca^2+^ signals in intact pulmonary endothelium was recorded in an *en face* preparation of small pulmonary arteries (PAs; ∼50 μm diameter with single smooth muscle cell layer (*21*); Fig. 1B) loaded with Fluo-4 AM (10 μM) using high-speed spinning-disk confocal Ca^2+^ imaging (Fig. 1B). Baseline Ca^2+^ signaling activity (number of Ca^2+^ event sites per EC and number of Ca^2+^ events per EC) was higher in the endothelium from mice exposed to IR compared with that in sham mice (Fig. 1C, left, Fig. S1). Our previous studies showed that baseline activity of Ca^2+^ signals in the PA endothelium primarily consists of inositol triphosphate receptor (IP_3_R)-mediated Ca^2+^ release from intracellular stores and Ca^2+^ influx through TRPV4 channels (*10, 11, 22*). The specific TRPV4 channel inhibitor, GSK2193874 (100 nM; hereafter, GSK219), completely abolished the increase in baseline endothelial Ca^2+^ signals after lung IR in control mice (Fig. 1D). Moreover, the lung IR-induced increase in endothelial Ca^2+^ signaling activity was lost in small PAs from TRPV4_EC_^-/-^ mice (Fig. 1C and 1D). These results suggest that increased TRPV4_EC_ channel activity mediates IR-induced increases in endothelial Ca^2+^ signals. To verify this possibility, we studied TRPV4_EC_ channel activity in the form of TRPV4_EC_ sparklets (*23*) (individual Ca^2+^-influx signals through TRPV4_EC_ channels) in the intact endothelium from small PAs of mice that underwent lung IR or sham surgery. TRPV4_EC_ sparklets were recorded in the presence of 20 μM cyclopiazonic acid (CPA; sarco/endoplasmic reticulum ATPase [SERCA] inhibitor) to eliminate interference from intracellular Ca^2+^-release signals. CPA, by itself, does not alter the activity of TRPV4 channels (*19, 24*). TRPV4_EC_ sparklet activity, both sparklet sites per cell and activity per site, was increased in mice exposed to IR compared with those receiving sham surgery (Fig. 1E and 1F). The IR-induced increase in TRPV4_EC_ sparklet activity was inhibited by the TRPV4 inhibitor GSK219 and was absent in the endothelium from TRPV4_EC-_^/-^ mice (Fig. 1E and 1F). In small PAs from mice that underwent 1-hour of left lung ischemia but not reperfusion, endothelial Ca^2+^ signaling activity was not altered (Fig. S2), suggesting that reperfusion was required for IR-induced increase in pulmonary endothelial Ca^2+^ signaling. These results provide the first direct evidence that lung IR increases the activity of TRPV4_EC_ channels in the pulmonary endothelium.

**Figure 1.**
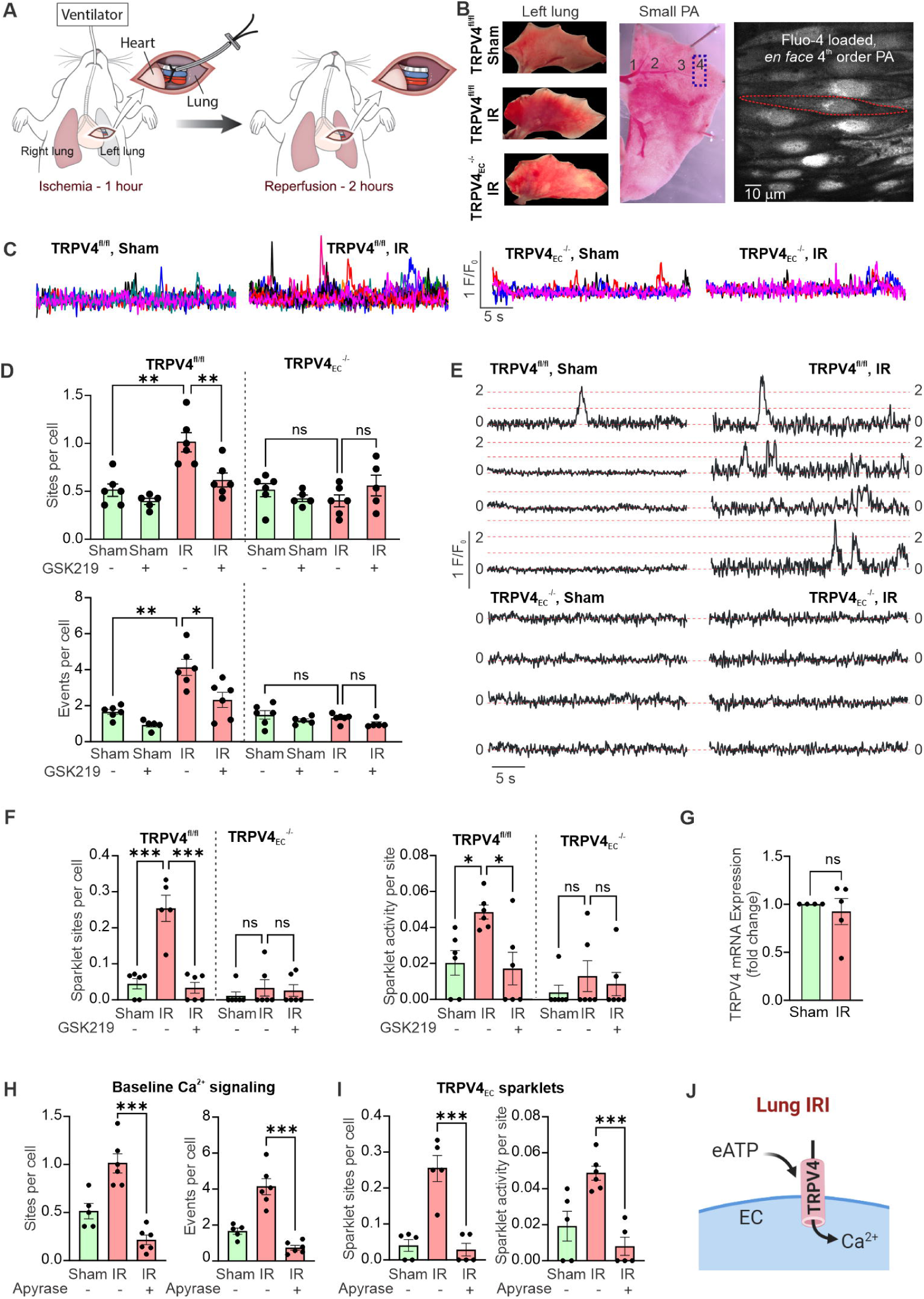
Lung IR increases TRPV4 channel activity in the pulmonary endothelium. (A) Left-lung hilar-ligation model of IRI (see Methods). Mice in the IR group underwent left lung hilar ligation for 1 hour followed by reperfusion for 2 hours. Control mice underwent sham surgery with no hilar ligation. (B) ***Left*,** representative images of left lungs from TRPV4^fl/fl^ sham, TRPV4^fl/fl^ IR, and TRPV4_EC_^-/-^ IR mice showing gross injury after IR and protection in TRPV4_EC_^-/-^ mice. ***Center,*** image showing the ordering system used for dissecting out small PAs (4^th^-order, dashed area) from left lungs for analysis of overall Ca^2+^ signals and TRPV4 sparklet activity. ***Right,*** grayscale image showing approximately 15 ECs from a field of view in a PA loaded with Fluo-4 AM. Red outline shows one EC. (C) Representative fractional fluorescence (F/F_0_) traces showing baseline Ca^2+^ signaling activity in the pulmonary endothelium from TRPV4^fl/fl^ sham, TRPV4^fl/fl^ IR, TRPV4_EC_^-/-^ sham, and TRPV4_EC_^-/-^ IR mice. (D) Total baseline Ca^2+^ signaling activity, represented as sites per cell (top) and events per cell (bottom), in PAs from TRPV4^fl/fl^ sham, TRPV4^fl/fl^ IR, TRPV4_EC_^-/-^ sham, and TRPV4_EC_^-/-^ IR mice, with and without the TRPV4 inhibitor GSK219 (100 nM) (n = 5–6/group, *P < 0.05, **P < 0.01; ns, not significant; two-way ANOVA). (E) Representative F/F_0_ traces reflecting TRPV4 sparklet activity in the presence of cyclopiazonic acid (CPA, 20 μM), used to eliminate intracellular Ca^2+^ release signals, in the pulmonary endothelium from TRPV4^fl/fl^ sham, TRPV4^fl/fl^ IR, TRPV4_EC_^-/-^ sham, and TRPV4_EC_^-/-^ IR mice. Dotted lines represent quantal levels (single-channel amplitudes) determined from all-point histograms.(*11*) (F) TRPV4 sparklet activity per site (**left**; NP_O_ per site, where N is the number of channels and P_O_ is the open-state probability) and TRPV4 sparklet sites per cell (**right**) in the presence of CPA in the pulmonary endothelium from TRPV4^fl/fl^ sham, TRPV4^fl/fl^ IR, TRPV4_EC_^-/-^ sham, and TRPV4_EC_^-/-^ IR mice (n = 5–6/group; *P < 0.05, ***P < 0.001; ns, not significant; two-way ANOVA). (G) TRPV4 mRNA levels in freshly isolated pulmonary endothelium from TRPV4^fl/fl^ mice, expressed relative to sham (n = 4) and IR (n = 5) (ns, not significant; unpaired t-test). (H) Baseline endothelial Ca^2+^ signaling activity in small PAs from TRPV4^fl/fl^ mice in the presence of apyrase (10 U/mL), an eATP-diphosphohydrolase, presented as sites per cell **(left)** and events per cell **(right)** (n = 5–6/group; ***P < 0.001; two-way ANOVA). (I) Endothelial TRPV4 sparklet sites per cell and sparklet activity in small PAs from TRPV4^fl/fl^ mice in the presence of CPA in pulmonary endothelium from TRPV4^fl/fl^ sham, TRPV4^fl/fl^ IR, and TRPV4_EC_^-/-^ IR mice (n = 5–6/group; ***P < 0.001; two-way ANOVA). (J) Diagram depicting induction of TRPV4 Ca^2+^ channel activity by eATP after lung IR, leading to elevated intracellular Ca^2+^.

Next, we tested the possibility that increased expression of TRPV4_EC_ channels underlies the increase in channel activity after IR. A quantitative polymerase chain reaction (qPCR) analysis of freshly isolated ECs from small PAs revealed that TRPV4 mRNA levels were not different between lung IR and sham groups (Fig. 1G), implying that increased TRPV4_EC_ channel activity after lung IR is not attributable to increased channel expression. In this regard, recent studies suggest that eATP is an endogenous activator of TRPV4_EC_ channels in the intact pulmonary endothelium (*8*). Hydrolyzing eATP with apyrase (10 U/mL) abolished IR-induced increases in baseline Ca^2+^ signaling activity and TRPV4_EC_ channel activity in small PAs (Fig. 1H and 1I), identifying eATP as a novel mediator of increased TRPV4_EC_ channel activity after lung IR (Fig. 1J).

### Endothelial P2Y2R signaling activates TRPV4 channel-dependent Ca^2+^ influx in lung IR

We earlier showed that, under healthy conditions, eATP activates TRPV4_EC_ channels through purinergic P2Y2R signaling in small PAs (*8*). Therefore, we used the P2Y2R inhibitor, AR-C 118925XX (ARC; 10 μmol*/*L), to test the role of P2Y2Rs in lung IR-induced increases in TRPV4_EC_ channel activity in small PAs. Treatment with ARC abolished lung IR-induced increases in baseline endothelial Ca^2+^ signaling activity in small PAs (Fig. 2A). ARC also inhibited lung IR-induced increases in TRPV4_EC_ sparklet activity (Fig. 2B). To test whether endothelial P2Y2Rs mediate TRPV4_EC_ channel activation after lung IR, we used tamoxifen-inducible, endothelium-specific P2Y2R- knockout (P2Y2R_EC_^-/-^) mice (*8*). P2Y2R_EC_^-/-^ and control (P2Y2R^fl/fl^) mice underwent lung IR or sham surgeries. IR-induced gross injury was apparent in left lungs from P2Y2R^fl/fl^ mice, but not in P2Y2R_EC_^-/-^ mice (Fig. 2C). Baseline Ca^2+^ signaling activity (Fig. 2D and 2E) and TRPV4_EC_ sparklet activity (Fig. 2F and 2G) were increased in control mice after IR compared with those after sham surgery. IR-induced increases in baseline Ca^2+^ signaling activity and TRPV4_EC_ sparklet activity were absent in the endothelium from P2Y2R_EC_^-/-^ mice (Fig. 2E–G). Moreover, lung IR did not alter the expression of endothelial P2Y2Rs at the mRNA level (Fig. 2H), implying that lung IRI involves increased activation, not upregulation, of endothelial P2Y2Rs. These data provide strong evidence that eATP acts through P2Y2R_EC_ to activate TRPV4_EC_ channels in lung IRI (Fig. 1I).

**Figure 2.**
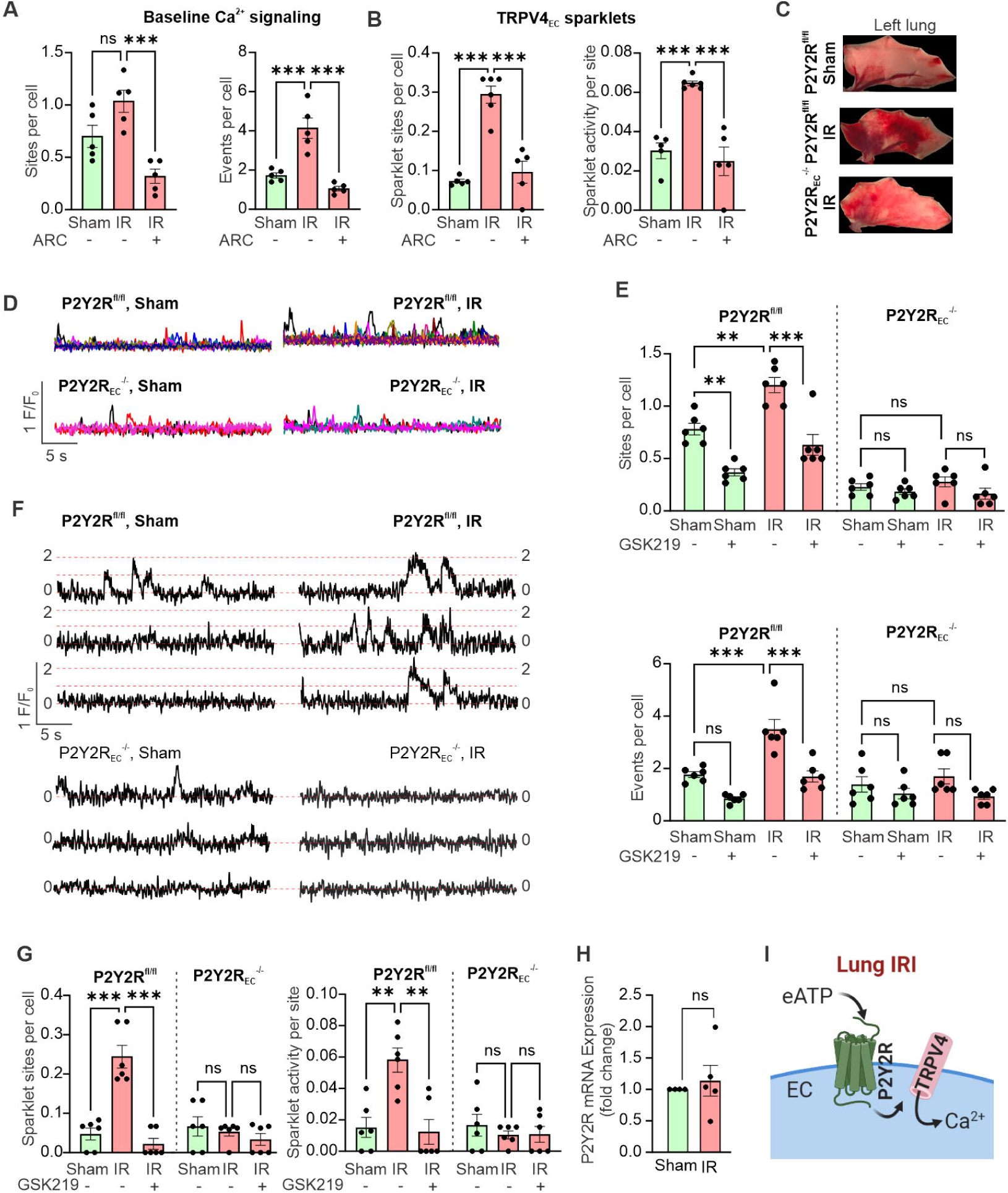
Lung IR increases TRPV4 sparklet activity via P2Y2R signaling in the pulmonary endothelium. Ca^2+^ signals were recorded from the endothelium of Fluo-4- loaded small PAs. (A) Total Ca^2+^ signaling activity (sites per cell and events per cell) and (B) TRPV4 sparklet activity (sparklet sites per cell and sparklet activity per site) in ECs from small PAs of C57BL6/J mice in the absence and presence of the P2Y2R inhibitor, AR-C 118925XX (ARC, 10 μM) (n = 5–6/group; ***P < 0.001; ns, not significant; two-way ANOVA). (C) Representative images of left lungs from P2Y2R^fl/fl^ sham, P2Y2R^fl/fl^ IR, and P2Y2R_EC_^-/-^ IR mice showing gross lung injury after IR and protection in P2Y2R_EC_^-/-^ mice. (D) Representative F/F_0_ traces showing baseline Ca^2+^ signaling activity in the pulmonary endothelium. (E) Dot plot showing total baseline Ca^2+^ signaling activity for P2Y2R^fl/fl^ sham, P2Y2R^fl/fl^ IR, P2Y2R_EC_^-/-^ sham, and P2Y2R_EC_^-/-^ IR mice, with and without the TRPV4 inhibitor GSK219 (100 nM) (n = 5–6/group; **P < 0.01, ***P < 0.001; ns, not significant; two-way ANOVA). (F) Representative TRPV4 sparklet traces and (G) analysis of sparklet activity in the pulmonary endothelium from P2Y2R^fl/fl^ sham, P2Y2R^fl/fl^ IR, P2Y2R_EC_^-/-^ sham, and P2Y2R_EC_^-/-^ IR mice (n = 5–6/group; **P < 0.01, ***P < 0.001; ns, not significant; two-way ANOVA). (H) TRPV4 mRNA levels in freshly isolated pulmonary ECs from P2Y2R^fl/fl^ mice (sham, n = 4; IR, n = 5; unpaired t-test). Data are expressed relative to the sham group. (I) Diagram depicting eATP-induced activation of endothelial P2Y2R signaling and Ca^2+^ influx through TRPV4 channels after lung IR.

### Endothelial P2Y2R deletion attenuates lung IRI

Compared with sham mice, control P2Y2R^fl/fl^ mice exposed to IR developed significant pulmonary dysfunction, as measured by reduced partial pressure of oxygen (PaO_2_) and lung compliance (Fig. 3A and 3B). P2Y2R_EC_^-/-^ mice undergoing IR exhibited significantly attenuated lung dysfunction after IR, as demonstrated by improved PaO_2_ and compliance (Fig. 3A and 3B). Significant pulmonary edema, as measured by lung wet-to-dry weight ratio, occurred in P2Y2R^fl/fl^ mice after IR, which was prevented in P2Y2R_EC_^-/-^ mice (Fig. 3C), suggesting preservation of the vascular endothelial barrier. The lung endothelial barrier was also assessed by measuring neutrophil infiltration (which requires migration across the endothelial barrier) and pro-inflammatory cytokine levels in BAL fluid. Neutrophil infiltration in lung tissue was markedly increased in control P2Y2R^fl/fl^ mice after IR (Fig. 3D), consistent with impaired endothelial barrier function. IR-induced neutrophil infiltration was significantly reduced in lungs from P2Y2R_EC_^-/-^ mice (Fig. 3D), providing evidence that endothelial P2Y2R deletion preserves endothelial barrier function after lung IR. Moreover, P2Y2R^fl/fl^ mice undergoing IR exhibited significantly higher levels of proinflammatory cytokines, including chemokine (C-X-C motif) ligand 1 (CXCL1/KC), chemokine (C-X-C motif) ligand 2 (CXCL2/MCP-1), interleukin 6 (IL-6), and tumor necrosis factor alpha (TNFα) (Fig. 3E). Expression of these cytokines was significantly reduced in P2Y2R_EC_^-/-^ mice after lung IR. Collectively, these results supported a crucial role for endothelial P2Y2Rs in lung IR-induced pulmonary dysfunction, edema, inflammation, and endothelial barrier disruption.

**Figure 3.**
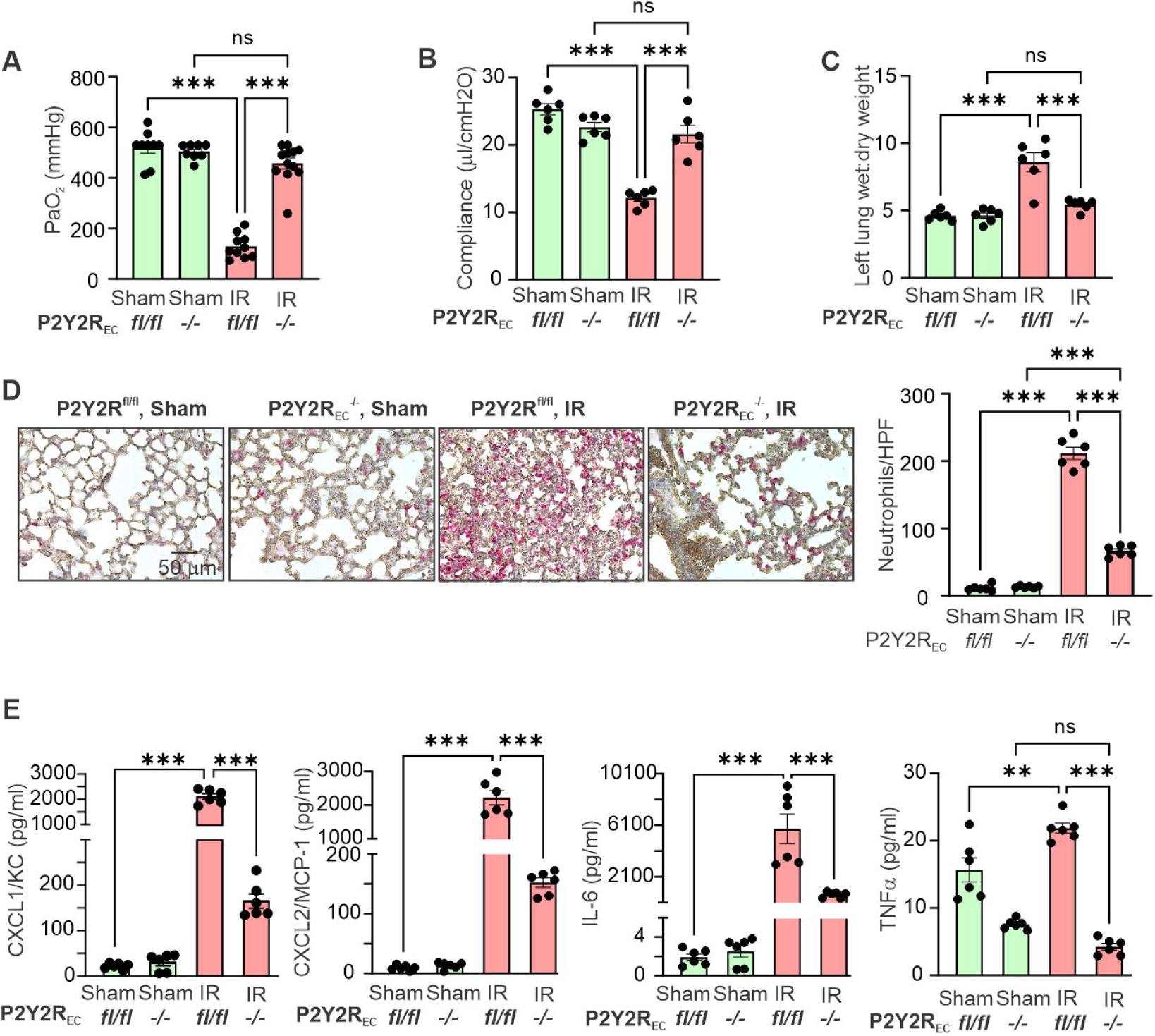
Endothelium-specific P2Y2R deletion decreases IR-induced neutrophil infiltration and lung edema and improves lung function after IR. (A) Partial pressure of arterial oxygen (PaO_2_), (B) lung compliance, and (C) lung edema (wet/dry weight ratio), measured in P2Y2R^fl/fl^ sham, P2Y2R^fl/fl^ IR, P2Y2R_EC_^-/-^ sham, and P2Y2R_EC_^-/-^ IR mice (n = 5–6/group; ***P < 0.001; ns, not significant; two-way ANOVA). (D) Representative immunostaining images showing neutrophil infiltration (stained pink with alkaline phosphatase). Scale bar: 50 μm. Quantification of the number of neutrophils per high- powered field (HPF; right) from P2Y2R^fl/fl^ sham, P2Y2R^fl/fl^ IR, P2Y2R_EC_^-/-^ sham, and P2Y2R_EC_^-/-^ IR mice (n = 5–6/group; ***P < 0.001; two-way ANOVA). (E) Concentrations of the proinflammatory cytokines CXCL1/KC, CXCL2/MCP-1, IL-6, and TNFa in left-lung BAL fluid from P2Y2R^fl/fl^ sham, P2Y2R^fl/fl^ IR, P2Y2R_EC_^-/-^ sham, and P2Y2R_EC_^-/-^ IR mice (n = 5–6/group; ***P < 0.001; ns, not significant; two-way ANOVA).

### Acute hypoxia-reoxygenation (HR) induces P2Y2R–TRPV4 signaling in the pulmonary endothelium

To determine whether IR-induced increases in P2Y2R_EC_–TRPV4_EC_ signaling are a result of hypoxia (ischemia) followed by reoxygenation (reperfusion), we used a novel *ex vivo* surrogate model (Fig. 4A) of IR, in which small PAs from TRPV4_EC_^-/-^, P2Y2R_EC_^-/-^, and respective control mice are exposed to 1-hour hypoxia followed by 10 minutes of reoxygenation (HR) or to normoxia for 70 minutes. The Ca^2+^ indicator, Fluo-4 AM (10 μM), was included during the final 10 minutes of incubation for each group. Acute HR exposure increased the activity of baseline endothelial Ca^2+^ signaling (Fig. 4B) and TRPV4 sparklets (Fig. 4C) in small PAs from TRPV4^fl/fl^ mice compared with that observed following exposure to normoxia. Small PAs from TRPV4_EC_^-/-^ mice were protected against HR-induced increases in baseline endothelial Ca^2+^ signaling and TRPV4_EC_ sparklet activity. Moreover, treatment with the TRPV4 inhibitor, GSK219 (100 nM), completely abolished the increase in endothelial Ca^2+^ signals in small PAs from TRPV4^fl/fl^ mice exposed to HR (Fig. 4B and 4C). Similar to the case for the lung IR model, HR-induced increases in endothelial Ca^2+^ signaling (Fig. 4D) and TRPV4 sparklet (Fig. 4E) activity were absent in small PAs from P2Y2R_EC_^-/-^ mice. HR also increased baseline Ca^2+^ signaling and TRPV4 sparklets *in vitro* in murine primary pulmonary microvascular endothelial cells (PMVECs), in association with an increase in ionic currents through TRPV4 channels (Fig. S3), indicating that direct exposure to acute HR increases TRPV4_EC_ activity in small PAs as well as PMVECs. Hypoxia alone (induced using sodium dithionite, 1 mM) (*25*) did not alter baseline endothelial Ca^2+^ signaling activity in small PAs (Fig. S4), suggesting that reoxygenation was required for HR-induced increase in pulmonary endothelial Ca^2+^ signals. Collectively, these results suggest that acute HR exposure increases TRPV4_EC_ channel activity in the pulmonary endothelium through P2Y2R signaling.

**Figure 4.**
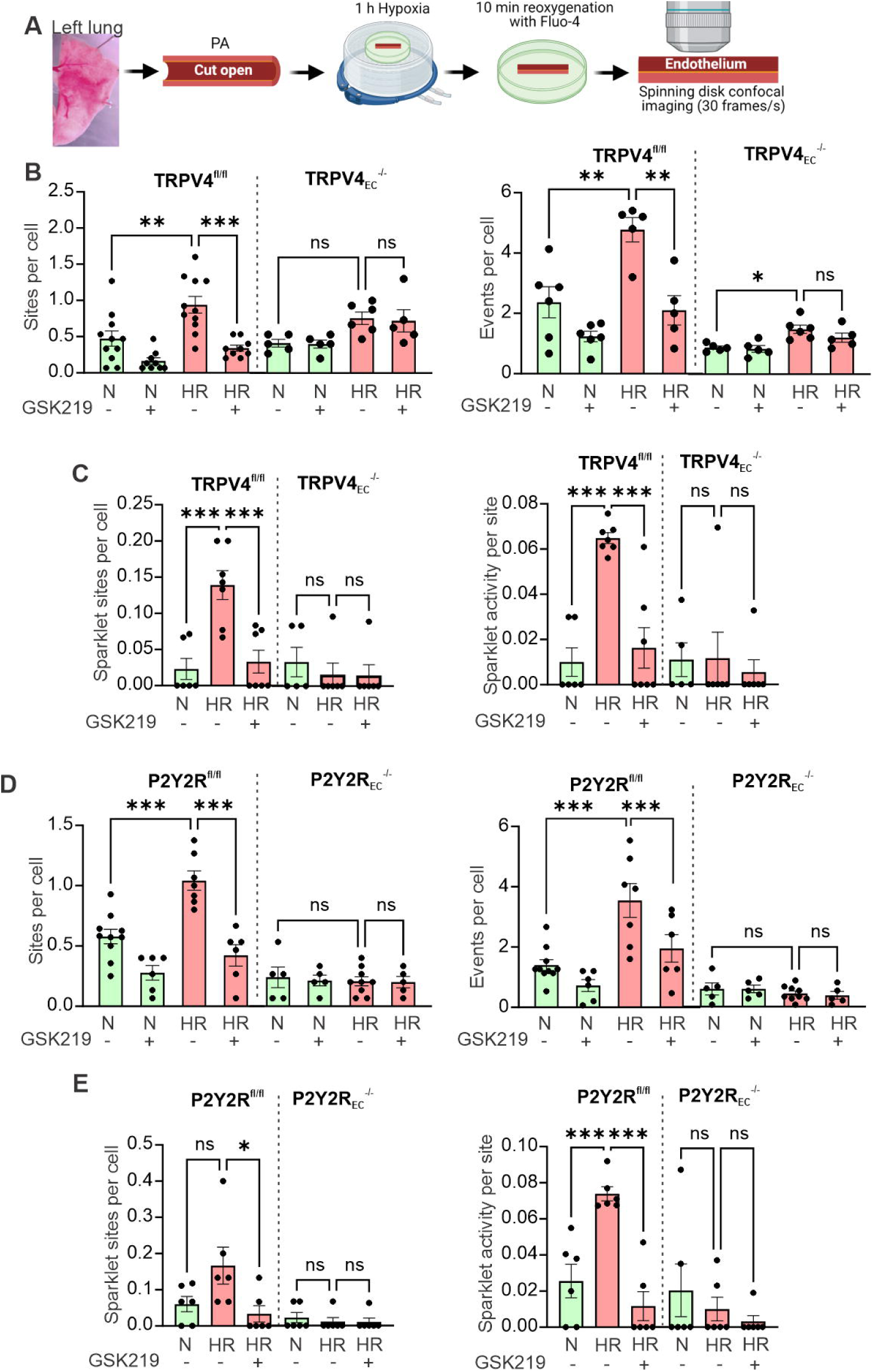
Exposure of small PAs to acute HR induces endothelial TRPV4 sparklet activity through P2Y2R stimulation. *En face* small PAs were exposed to 1 hour hypoxia followed by 10 minutes reoxygenation (HR) or normoxia (N) for 70 minutes. The final 10 minutes included incubation with the Ca^2+^ indicator, Fluo-4 (10 μM), for each group. (A) HR model and (B) analysis of baseline Ca^2+^ signaling activity and (C) TRPV4 sparklet activity after normoxia or HR in the pulmonary endothelium of small PAs from TRPV4^fl/fl^ and TRPV4_EC_^-/-^ mice, in the presence or absence of the TRPV4 inhibitor, GSK219 (100 nM) (n = 5–6/group; ***P < 0.001; ns, not significant; two-way ANOVA). (D) Total baseline Ca^2+^ signaling activity and (E) TRPV4 sparklet activity after normoxia or HR in the pulmonary endothelium of small PAs from P2Y2R^fl/fl^ and P2Y2R_EC_^-/-^ IR mice, in the presence and absence of the TRPV4 inhibitor GSK219 (n = 5–6/group; *P < 0.05, ***P < 0.001; ns, not significant; two-way ANOVA).

### Acute HR exposure increases TRPV4 channel activity in a P2Y2R-dependent manner in human pulmonary ECs

To determine whether acute HR-induced increase in P2Y2R–TRPV4 signaling is also observed in human pulmonary endothelium, we used freshly isolated small PAs (∼ 50 μm) from normal-looking wedge samples (Fig. 5A) of human lungs obtained during lung transplant surgeries. Human small PAs were exposed to HR (1 hour hypoxia followed by 10 minutes reoxygenation), after which endothelial Ca^2+^ signals were recorded. Human small PAs also showed an increase in baseline endothelial Ca^2+^ signaling after HR exposure (Fig. 5B and 5C). Moreover, HR-induced increase in endothelial Ca^2+^ signaling was inhibited by TRPV4 inhibitor GSK219 (Fig. 5C) and P2Y2R inhibitor ARC (Fig. 5D) in human small PAs. These data provided first evidence that HR exposure results in a P2Y2R- and TRPV4-dependent increase in endothelial Ca^2+^ signaling in human pulmonary microvascular endothelium.

**Figure 5.**
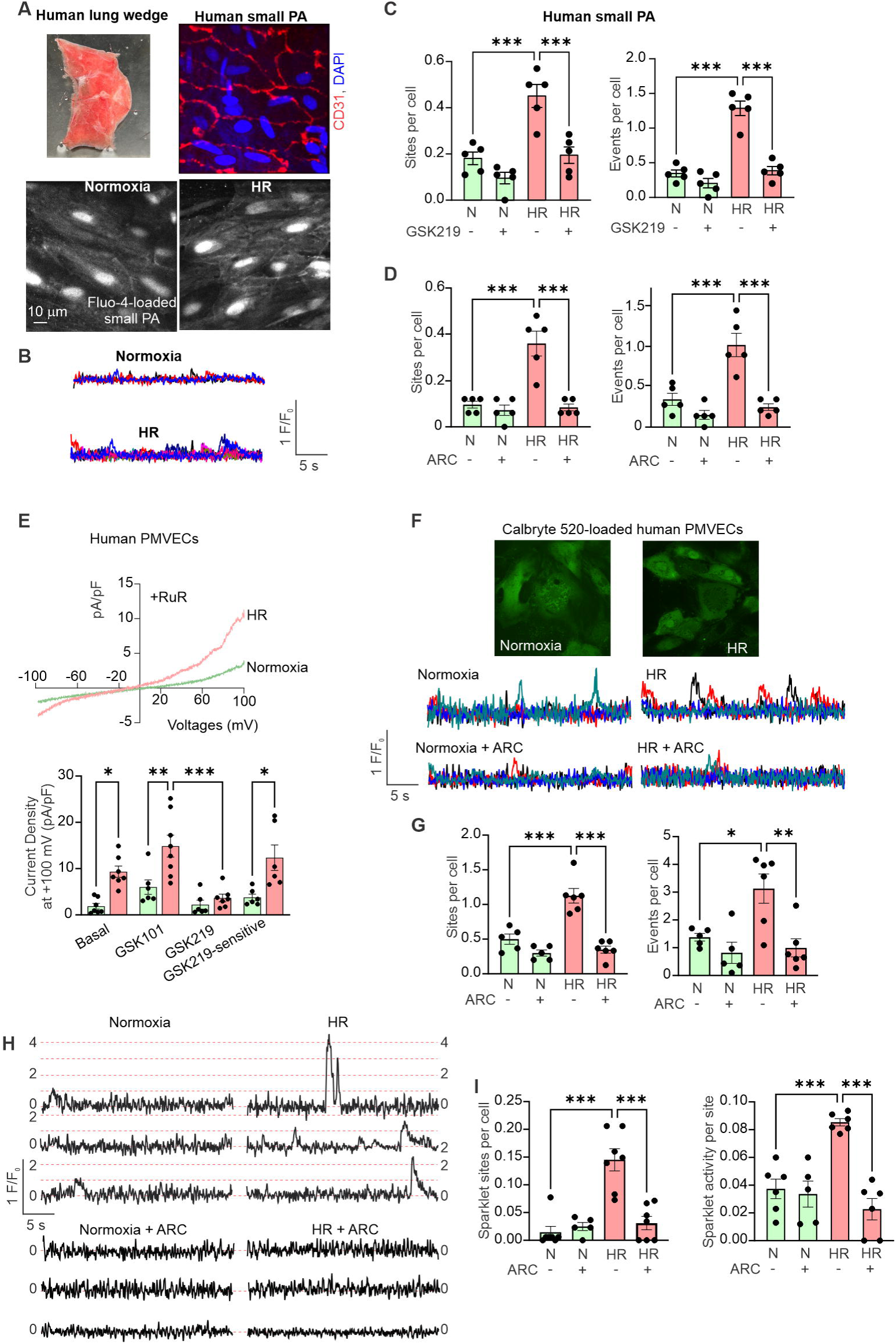
Exposure to acute HR increases TRPV4 channel activity in human PMVECs. (A) ***Top left*,** an image showing a lung wedge sample obtained during lung transplantation surgery. ***Top right*,** CD31 immunostaining showing ECs in small PAs isolated from human lungs (left). ***Bottom*,** greyscale images showing ECs from fluo-4- loaded, freshly isolated small PAs from human lungs. Small PAs were exposed to normoxia or HR. (B) F/F_0_ traces showing baseline endothelial Ca^2+^ signaling activity in small PAs from human lungs (right). (C) Analyzed data showing total baseline endothelial Ca^2+^ signaling activity in the absence and presence of GSK219 (100 nM) in small PAs from human lungs (n = 5/group, ***P < 0.001, two-way ANOVA). (D) Analyzed data showing total baseline endothelial Ca^2+^ signaling activity in the absence and presence of ARC (10 μM) in small PAs from human lungs (n = 5/group, ***P < 0.001, two-way ANOVA). (E) ***Top*,** representative whole-cell patch-clamp traces of TRPV4 currents, defined as currents in the presence of the TRPV4 agonist GSK101 (10 nM) minus those in the presence of GSK101 + the TRPV4 antagonist GSK219 (100 nM), in human PMVECs. Outward currents through TRPV4 channels were recorded using −100 mV to +100 mV voltage ramps. Experiments were performed in the presence of ruthenium red (RuR; 1 µM) to prevent Ca^2+^ influx through TRPV4 channels and subsequent Ca^2+^- dependent activation of K^+^ channels. ***Bottom*,** averaged outward currents in human PMVECs at +100 mV under basal conditions, in the presence of GSK101 (10 nM) or GSK101 + GSK219 (100 nM), and GSK219-sensitive currents (GSK101 minus [GSK101 + GSK219]) (*P < 0.05, **P < 0.01, ***P < 0.001 vs. Basal; two-way ANOVA). (F) Calbryte 520-loaded human PMVECs (top) and representative traces (bottom) and analyzed data (G) showing total baseline Ca^2+^ signaling activity in the absence and presence of ARC (10 μM) in human PMVECs exposed to normoxia or HR. (H) Representative traces and (I) analyzed data showing TRPV4 sparklet activity in the absence and presence of ARC (10 μM) in human PMVECs (n = 5–7/group, ***P < 0.001; two-way ANOVA).

Next, we exposed human PMVECs to acute HR (1 hour hypoxia followed by 10 minutes reoxygenation) or to normoxia for 70 minutes, after which TRPV4_EC_ channel activity and endothelial Ca^2+^ signals were measured. TRPV4 currents, defined as those sensitive to the TRPV4 inhibitor, GSK219 (100 nM), were increased in human PMVECs (Fig. 5E) by HR exposure, suggesting that acute HR exposure increases TRPV4 channel activity in the human pulmonary endothelium.

To record individual Ca^2+^ signals in human PMVECs, we included the Ca^2+^ indicator, Calbryte520 (0.1 μM), during the last 10 minutes of incubation under reoxygenation or normoxia conditions. Baseline Ca^2+^ signaling (Fig. 5F and 5G) and TRPV4_EC_ sparklet activity (Fig. 5H and 5I) were increased in human PMVECs after HR exposure. Similar to our findings in murine PMVECs, HR induced increase in baseline endothelial Ca^2+^ signaling as well as TRPV4_EC_ sparklets in human PMVECs was blocked by treatment with the selective P2Y2R inhibitor, ARC (Fig. 5F-5I). These findings indicate that acute HR exposure induces TRPV4_EC_ channel activity in a P2Y2R-dependent manner in both mouse and human PMVECs.

### eATP efflux through endothelial Panx1 channels mediates IR-induced activation of P2Y2R_EC_–TRPV4_EC_ signaling in the pulmonary endothelium

We have recently shown that activation of endothelial Panx1 channels increases eATP levels in small PAs under normal conditions (*8*). Moreover, lung IR increased eATP levels in BAL fluid through endothelial Panx1 activation (*9*). Consistent with these reports, we show that Panx1_EC_^-/-^ mice are protected against IR-induced gross lung injury (Fig. 6A). Bioluminescence measurements showed that lung IR induced eATP release from small PAs; this effect was absent in small PAs from Panx1_EC_^-/-^ mice (Fig. 6B), suggesting that lung IR increases eATP efflux through endothelial Panx1 channels. Endothelial Panx1 mRNA expression was not altered after lung IR (Fig. 6C), suggesting that increased eATP efflux after lung IR is a result of increased activity, not expression, of endothelial Panx1 channels. To determine the effect of Panx1-mediated eATP efflux on endothelial Ca^2+^ signaling after lung IR, we recorded baseline endothelial Ca^2+^ signals and TRPV4 sparklet activity in small PAs from Panx1_EC_^-/-^ and Panx1^fl/fl^ mice after IR or sham surgery. As expected, the activity of baseline endothelial Ca^2+^ signaling (Fig. 6D) and TRPV4_EC_ sparklets (Fig. 6E) was elevated in control Panx1^fl/fl^ mice after lung IR compared with sham mice. However, no increase in baseline endothelial Ca^2+^ signaling or TRPV4_EC_ sparklets was observed in the endothelium from Panx1_EC_^-/-^ mice following IR (Fig. 6D and 6E). Taken together, these data indicate that increased eATP efflux through endothelial Panx1 channels activates downstream P2Y2R_EC_–TRPV4_EC_ signaling after lung IR. In addition, HR-induced increases in the baseline activity of endothelial Ca^2+^ signals (Fig. 6F) and TRPV4_EC_ sparklets (Fig. 6G) were also absent in *ex vivo* small PAs from Panx1_EC_^-/-^ mice. Collectively, these findings support the concept that eATP effluxed via endothelial Panx1 channels activates endothelial P2Y2R–TRPV4 signaling, resulting in lung inflammation, endothelial barrier disruption, edema, and lung dysfunction after lung IR (Fig. 7).

**Figure 6.**
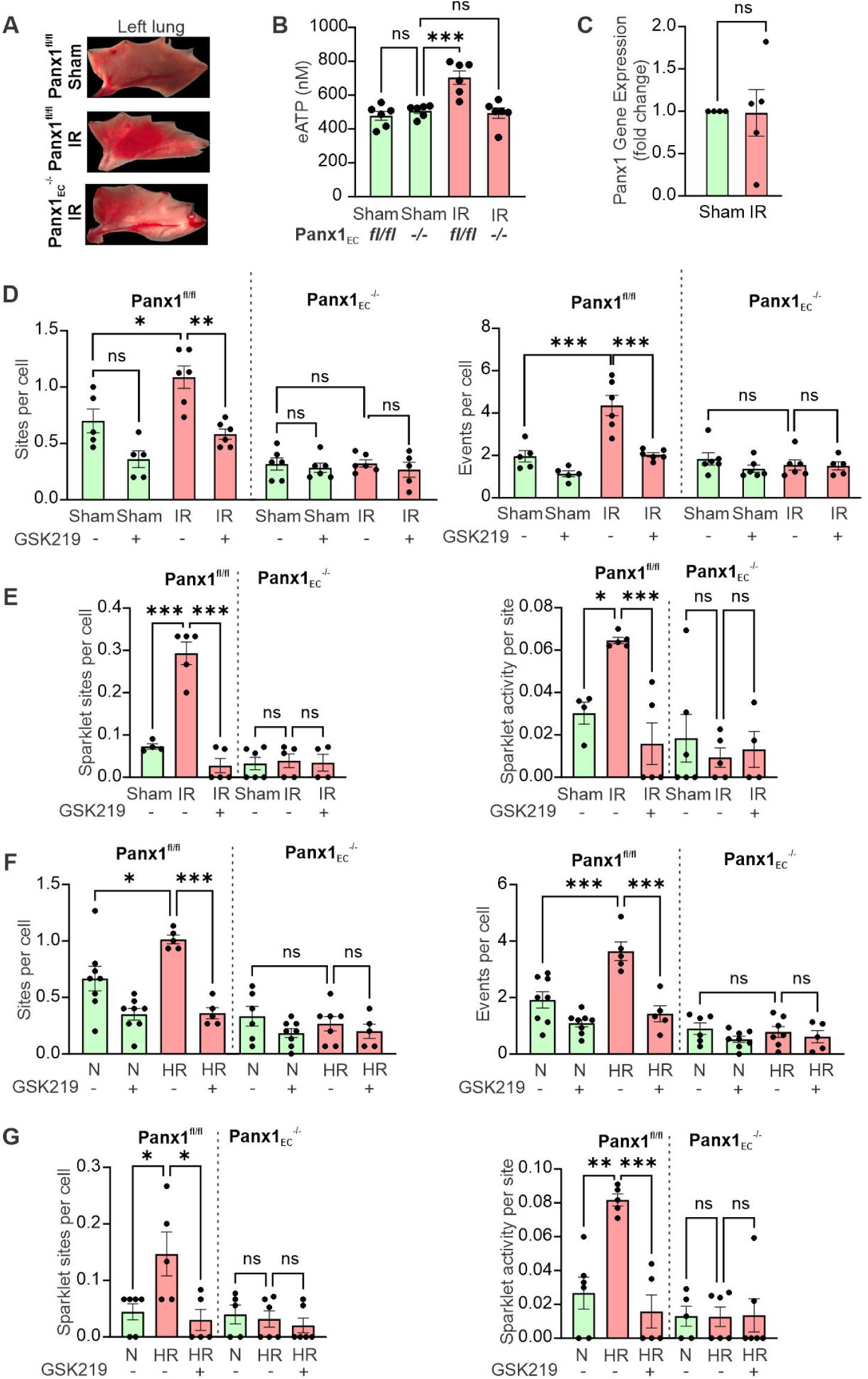
Panx1-effluxed eATP mediates IR-induced activation of endothelial P2Y2R–TRPV4 signaling. (A) Representative images of left lungs from Panx1^fl/fl^ sham, Panx1^fl/fl^ IR, and Panx1_EC_^-/-^ IR mice showing gross injury after IR and protection in Panx1_EC_^-/-^ mice. (B) Bioluminescence measurements of eATP (nM) released from small PAs of Panx1^fl/fl^ sham, Panx1^fl/fl^ IR, Panx1_EC_^-/-^ sham, and Panx1_EC_^-/-^ IR mice (n = 6/group; ***P < 0.001; ns, not significant; two-way ANOVA). (C) Panx1 mRNA levels in freshly isolated pulmonary ECs from Panx1^fl/fl^ sham (n = 4) and Panx1_EC_^fl/fl^ IR (n = 5) mice (ns, not significant; unpaired t-test). Data are expressed relative to the sham group. (D) Total baseline Ca^2+^ signaling activity and (E) TRPV4 sparklet activity in the pulmonary endothelium from Panx1^fl/fl^ sham, Panx1^fl/fl^ IR, Panx1_EC_^-/-^ IR, and Panx1_EC_^-/-^ IR mice in the absence and presence of the TRPV4 inhibitor GSK219 (100 nM) (n = 5–6/group, *P < 0.05, **P < 0.01, ***P < 0.001; ns, not significant; two-way ANOVA). (F) Total baseline Ca^2+^ signaling activity and (G) TRPV4 sparklet activity in the pulmonary endothelium from Panx1^fl/fl^ sham and Panx1_EC_^-/-^ IR mice in the absence and presence of GSK219 (100 nM) (n = 5–6/group, *P < 0.05, *P < 0.01, ***P < 0.001; ns, not significant; two-way ANOVA).

**Figure 7.**
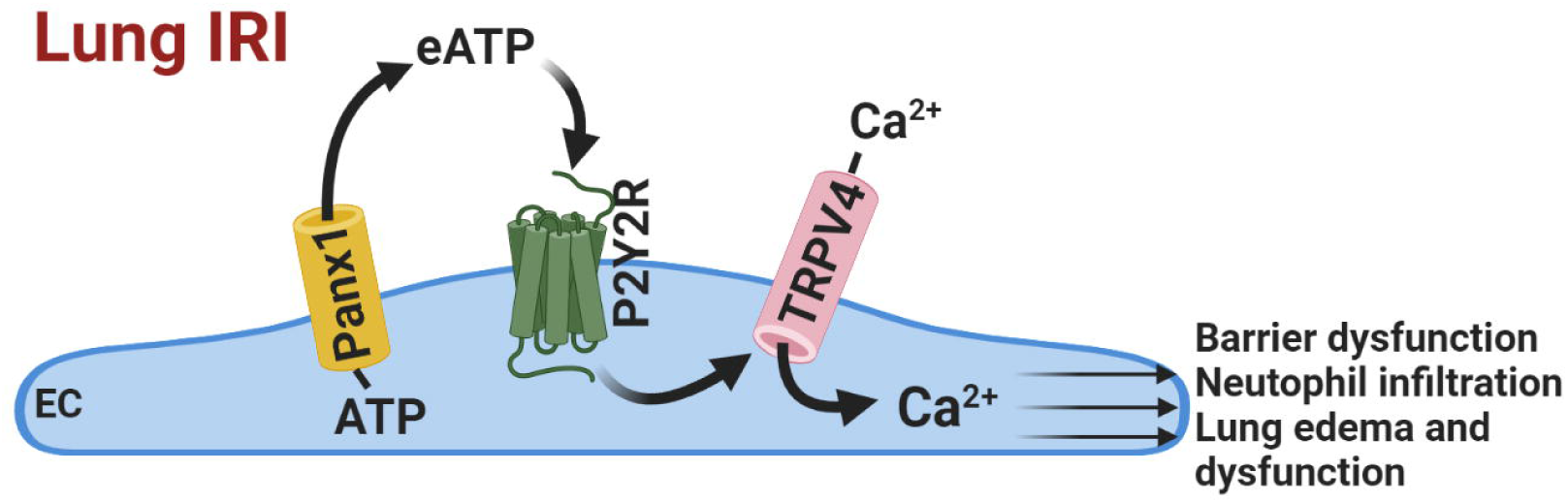
Outline of lung IR-induced Panx1–eATP–P2Y2R–TRPV4 signaling in pulmonary ECs. eATP, effluxed through endothelial Panx1 channels, activates the endothelial P2Y2R, which, in turn, increases Ca^2+^ influx via activation of endothelial TRPV4 channels to mediate endothelial barrier dysfunction, neutrophil infiltration into the alveolar space, lung edema, and dysfunction after lung IR.

## Discussion

Several previous studies provided evidence that endothelial TRPV4 channels are involved in various models of acute lung injury (*12, 13, 15*). We recently showed that endothelial TRPV4 channels play a critical role in lung IRI (*16*). Our current study provides novel insights into the purinergic mechanisms that mediate lung IR-induced activation of endothelial TRPV4 channels and subsequent lung injury. We show that lung IR increases endothelial Panx1 channel-mediated efflux of eATP, which then activates endothelial P2Y2R signaling. This, in turn, increases the activity of endothelial TRPV4 channels, resulting in endothelial barrier disruption, lung edema, immune cell infiltration, and lung dysfunction after IR. Activation of this endothelial Panx1–eATP–P2Y2R–TRPV4 signaling axis was observed in *in vivo*, *ex vivo*, and *in vitro* models of IR. Importantly, acute HR-induced P2Y2R–TRPV4 signaling was also observed in human PMVECs and endothelium from human small PAs. Overall, our data provide evidence for a novel role of endothelial P2Y2Rs in IR-induced lung edema and dysfunction and suggest that P2Y2Rs could be therapeutically targeted to prevent lung IRI, and subsequent primary graft dysfunction, after lung transplantation. The conclusions of this study may also assist in future investigations of the interplay between endothelial mechanisms underlying lung IRI with those from immune cells and epithelial cells.

eATP rapidly accumulates at sites of inflammation including IRI sites.(*9*) The proinflammatory effects of eATP are largely attributable to P1 and P2 receptors at the cell membrane. These receptors are widely expressed in pulmonary endothelium, epithelium, and a variety of inflammatory cells (*26–28*). G protein-coupled P2Y receptors (P2Y1,2,3,6,11-14) and ligand-gated P2X ion channels (P2X1-7) make up the P2 receptor family (*26, 29*). Among P2Y receptor subtypes, P2Y2R has been most commonly linked to inflammation and IRI (*26, 29*). P2Y2R deletion was shown to be protective in several models of acute inflammation (*26*). Moreover, endothelial P2Y2Rs were found to promote inflammatory cytokine production in atherosclerosis (*30*). P2X receptors, especially the P2X7 receptor, have also been shown to play important roles in inflammatory disorders (*26*). Our present study shows that the endothelial P2Y2R is critical for lung inflammation, edema, and dysfunction after lung IR. Further studies will be needed to define the role of P2X7R in endothelial/epithelial/immune cells, in lung IRI.

When activated by various stress stimuli, Panx1 channels on the cell membrane are known to release ATP into the extracellular space (*31*). Upstream mechanisms involved in increasing the activity of endothelial Panx1 channels in lung IRI are not yet known. Panx1 channels can be activated by many stimuli, including, but not limited to, TNFα, mechanical stretch, membrane depolarization, a rise in intracellular Ca^2+^, and temperature (*31, 32*). Notably, TNFα levels are increased in the lungs after IR, and we previously demonstrated that macrophage-dependent TNFα secretion is an early trigger for inflammation during lung IRI (*33, 34*). Our previous studies on the pulmonary endothelium showed that, under physiological conditions, Ca^2+^ influx through endothelial TRPV4 channels does not activate Panx1 (*8*). However, supraphysiological activation of TRPV4 channels with the agonist, GSK101 (10 nM), increased eATP efflux through Panx1 channels (Fig. S5), raising the interesting possibility that Panx1–P2Y2R–mediated stimulation of TRPV4 channels may lead to a feed-forward loop that activates Panx1, and in turn, results in further, rapid activation of TRPV4 channels.

Lung IRI involves a complex interplay between immune cells and pulmonary endothelial and epithelial cells (*16, 33, 35*). The contribution of Panx1, P2Y2R, and TRPV4 channels in epithelial and immune cells to lung IRI will be an interesting topic for future investigations. TRPV4 channels in epithelial cells and immune cells have been implicated in lung dysfunction and edema in lung injury (*36–38*). Panx1 expression has also been reported in epithelial and immune cells (*39, 40*). Our demonstration that IR-induced lung edema and dysfunction are absent in Panx1_EC_^-/-^(*9*), P2Y2R_EC_^-/-^, and TRPV4_EC-_^/-^(*16*) mice suggests a central role for an endothelial cell Panx1–P2Y2R–TRPV4 signaling axis in lung IRI. Studies in epithelial-, macrophage-, and neutrophil-specific Panx1/P2Y2R/TRPV4-knockout mice will be required to identify a potential role for a Panx1–P2Y2R–TRPV4 signaling axis in these cell types during lung IRI. Our results show that acute HR also increased the activity of the Panx1–P2Y2R–TRPV4 signaling axis in isolated small PAs as well as primary PMVECs. Because isolated small PAs and PMVECs (as used in our study) cannot recruit immune cells and these preparations lack epithelial cells, we posit that HR- or IR-induced effects on endothelial Ca^2+^ signaling originate at the level of pulmonary microvascular ECs.

The present study suggests that endothelial deletion of P2Y2R is sufficient to prevent IR-induced neutrophil infiltration. Functional P2Y2Rs are also expressed in neutrophils (*41*). However, the endothelial P2Y2R likely exerts indirect effects on neutrophil infiltration and function such that disruption of the pulmonary endothelial barrier by P2Y2R signaling increases neutrophil extravasation after IR. A similar mechanism was recently proposed to account for the beneficial effects of TRPV4 channel inhibition in a model of chemically induced lung injury (*15*). Our present study shows that IR-induced CXCL1, IL-6, and MCP-1 expression are significantly attenuated by endothelial P2Y2R deletion. We previously showed that endothelial deletion of TRPV4 attenuates IR-induced increases in several other inflammatory cytokines, including TNFα, IFNγ, and IL-17 (*5*). These inflammatory cytokines are largely produced by immune cells in models of lung IRI (*9, 35*). Our finding that endothelial deletion of P2Y2R reduces inflammatory cytokines in BAL fluid raises the interesting possibility that the endothelium may also be a source of inflammatory cytokines in lung IRI. Further studies are needed to investigate this possibility. Another possibility is that P2Y2R deletion from ECs preserves endothelial barrier after IR, thereby potentially dampening overall inflammation and activation of innate immune cells. Although endothelial P2Y2Rs appear to be essential for the production of inflammatory mediators during lung IRI, whether other cell types also contribute to this effect is not clear.

One of the limitations of the study is that the lung IRI model is not fully representative of clinical lung transplantation since there is no allograft transplantation involved. However, the IRI component of primary graft dysfunction can be modeled *in vivo* using the hilar-ligation approach, a well-accepted model that has been used to simulate post-transplant IRI (*42*). We previously showed that this model induces inflammatory pathways common to experimental lung transplant models (*9*). Additional studies using lung transplant models will be needed to confirm the effect of P2Y2R deletion/inhibition on lung IRI. Studies of individual Ca^2+^ signals in the intact endothelium of small PAs offers advantages over *in vitro* studies of EC monolayers or investigations of whole-cell increases in Ca^2+^. However, it is likely that most pathological events in lung IRI occur at the level of the pulmonary capillary endothelium or alveolo-capillary barrier in lung IRI. Since a reliable analysis of individual Ca^2+^ signals in the intact pulmonary capillary endothelium is not technically feasible, small PAs offer an appropriate alternative option for studying the effects of lung IR on endothelial Ca^2+^ signals. The small PAs used in the current study are ∼50 μm in diameter, have a single smooth muscle cell layer surrounding the EC layer (*21*), and lie close to the capillary bed.

In conclusion, our studies provide the first evidence that eATP, effluxed through endothelial Panx1 channels, activates endothelial P2Y2Rs, which, in turn, increases Ca^2+^ influx through TRPV4_EC_ channels to mediate endothelial barrier disruption, neutrophil infiltration, edema, and dysfunction after lung IR. Thus, this novel endothelial Panx1– P2Y2R–TRPV4 signaling axis may be a key pathway that could be targeted therapeutically to prevent primary graft dysfunction after lung transplantation.

## Materials and Methods

### Drugs and chemicals

Cyclopiazonic acid (CPA, depletes internal stores of Ca^2+^), GSK2193874 (GSK219, TRPV4 inhibitor), GSK1016790A (GSK101, TRPV4 agonist), AR-C 118925XX (ARC, P2Y2R inhibitor) and Ruthenium Red (TRPV4 channel blocker) were purchased from Tocris Bioscience (Minneapolis, MN). Fluo-4 AM (Ca^2+^ indicator) was purchased from Invitrogen (Carlsbad, CA). Tamoxifen and apyrase were obtained from Sigma-Aldrich (St. Louis, MO). Calbryte™ 520 AM (Ca^2+^ indicator) was purchased from AAT Bioquest (Pleasanton, CA). Pentobarbital was purchased from Sagent Pharmaceuticals (Schaumburg, IL). All other chemicals were purchased from Sigma-Aldrich.

### Animals

All animal protocols were approved by the University of Virginia Animal Care and Use Committee. All experiments were performed in 10-14 week old mice on a C57BL6/J background, in at least two independent batches. Both male and female mice were used. Lung IR-induced endothelial Ca^2+^ signaling and lung dysfunction were not different between male and female mice. TRPV4^fl/fl^ mice (*43*), P2Y2R^fl/fl^ mice (*17*), and Panx1^fl/fl^ mice (*44*) were crossed with Cdh5-CreERT2 mice (*45*) to generate EC-specific knockouts. EC-specific knockout of TRPV4, P2Y2R, and Panx1 was induced by feeding tamoxifen (40 mg/kg/d; Envigo Diet TD.130856) to 6-week-old TRPV4^fl/fl^ Cdh5-Cre, P2Y2R^fl/fl^ Cdh5-Cre, and Panx1^fl/fl^ Cdh5-Cre mice for 14 days. Mice were used for experiments after a 1-week tamoxifen washout period. Tamoxifen-fed TRPV4^fl/fl^, P2Y2R^fl/fl^, and Panx1^fl/fl^ mice were used as controls. Knockout validation information was provided previously (*8*). C57BL6/J mice were obtained from the Jackson Laboratory (Bar Harbor, ME). Mice were euthanized with pentobarbital (90 mg/kg, i.p.) followed by cranial cervical dislocation.

An independent team member performed random assignment of animals to groups and did not have knowledge of treatment assignment groups. All the experiments were performed in at least two independent batches. The experiments were blinded; information about the groups was withheld from the experimenter.

### Genotyping

Genomic DNA was extracted by treating samples of ear tissue with HotSHOT lysis buffer (25 mM NaOH, 0.2 mM EDTA) and then neutralizing with an equal volume of 40 mM Tris-HCL. PCR was performed on a T100 Thermal Cycler (Bio-Rad, Hercules, CA) using 1 unit of Bioline MangoTaq polymerase and buffer (London, England), 1.5 mM MgCl_2_, 200 µM each dNTP, 1 µM 5’ and 3’ primers, and ∼100–250 ng genomic DNA. Reaction products were run on a 1% agarose gel containing 0.2 µg/µL ethidium bromide in TAE buffer (40 mM Tris base, 20 mM acetic acid, 1 mM EDTA) at 90V. Gels were visualized using 302 nm UV light, and the sizes of PCR products were calibrated using a 100-bp DNA Ladder (New England BioLabs, Ipswich, MA). Mice were genotyped using the following primer pairs: Cdh5-Cre, 5’- GCC TGC ATT ACC GGT CGA TGC AAC GA-3’ (forward) and 5’-GTG GCA GAT GGC GCG GCA ACA CCA TT-3’ (reverse); TRPV4^fl/fl^, 5’-CAT GAA ATC TGA CCT CTT GTC CCC-3’ (forward) and 5’-CGG ACC ACA CGT CTG TCA TGT GTT-3’ (reverse); and P2Y2R^fl/fl^, 5’-TGA CGA CTC AAG ACG GAC AG-3 (forward) and 5’-TCC CAA CTC ACA CGT ACA AAT G-3’ (reverse). All primers were obtained from Eurofins Genomics (Louisville, KY). Panx1 genotyping (Panx1^fl/fl^) was performed by Transnetyx, Inc. (Memphis, TN) using the primer pair, 5’- CAA CAA GTT TGT ACA AA AAA GCA GGC TG-3’ (forward) and 5’- TGC TTG TAT AGA TAT CTG ACC CAG AGC T-3’ (reverse), and FAM reporter probe, CCC AGC TTT CTT GTA CAA.

### Murine lung IR model

A well-established left lung IRI model involving temporary *in vivo* ligation of the left hilum (pulmonary artery, vein, and airway) was used (*20*). Briefly, mice were anesthetized with inhaled isoflurane, intubated, ventilated with room air at 150 strokes/min with a tidal volume of 6 mL/g (Harvard Apparatus Co, South Natick, MA), and heparinized (20 U/kg; Hospira Inc, Lake Forest, IL). An anterolateral thoracotomy was performed through the fourth intercostal space to expose the left hilum. A 6-0 Prolene suture (Ethicon, Somerville, NJ) was passed around the left hilum, and the suture ends were passed through a 5-mm length of polyethylene (PE)-50 tubing. A surgical clip was used to fashion a Rummel tourniquet to provide temporary occlusion of the left hilar structures. The thoracotomy was closed, anesthesia was weaned, and analgesia was administered (intraperitoneal buprenorphine, 0.2 mg/kg). Animals recovered in their cages during the 1-hour ischemic period, after which they were anesthetized again and intubated, the Rummel tourniquet was removed, and the thoracotomy was closed. Animals were again extubated, recovered in their cage, and monitored during the 2-hour reperfusion period. Sham animals underwent the same surgical procedure except that application of the Rummel tourniquet to induce ischemia was omitted; thus, these animals were perfused for a total of 3 hours.

### Partial pressure of arterial oxygen (PaO_2_)

After reperfusion, animals were anesthetized, intubated, and ventilated with 100% oxygen. A clamshell thoracotomy was performed, and a 6-0 Prolene suture was passed around the right hilum. The right hilum was ligated, and the tidal volume was decreased to 4 mL/g. After 5 minutes of isolated left lung perfusion/ventilation, a left ventricular puncture was performed with a 25-gauge needle and blood was collected. Left lung-specific PaO_2_ was immediately measured using an i-STAT handheld blood analyzer and CG4+ cartridge (Abbott Point of Care, Princeton, NJ).

### Lung compliance

Total lung compliance was assessed in separate groups of animals using an IPL-1 isolated perfused lung system (Harvard Apparatus, Holliston, MA) as previously described (*16*). After reperfusion, animals were anesthetized with isoflurane, and a tracheostomy was performed. Animals were ventilated with room air at 150 strokes/min, a tidal volume of 200 mL, and a positive end-expiratory pressure of 2 cm H_2_O. Animals were exsanguinated by transaction of the inferior vena cava. The pulmonary artery was cannulated through the right ventricle, and a left ventricular incision was made toward the apex of the heart to facilitate venting. Lungs were perfused at 60 mL/g/min with 37°C Krebs-Henseleit buffer containing 0.20% glucose, 0.014% magnesium sulfate, 0.016% monopotassium phosphate, 0.035% potassium chloride, 0.69% sodium chloride, and 2.1% sodium carbonate. Ventilation and perfusion were maintained for a 5 minute equilibration period, and then lung compliance was recorded using the PULMODYN data-acquisition system (Hugo Sachs Elektronik, March-Hugstetten, Germany).

### Bronchoalveolar lavage (BAL) and cytokine analysis

Left lungs were lavaged with 0.4 mL cold phosphate-buffered saline (PBS). The resulting BAL fluid was centrifuged (1200 rpm) at 4°C for 5 minutes, and the supernatant was stored at −80°C. Cytokine concentrations in BAL fluid were measured using the multiplex Bio-Plex 200 System and Bio-Plex Pro Mouse Cytokine/Chemokine Immunoassays (Bio-Rad, Hercules, CA).

### Lung wet-to-dry weight ratio

Lung wet-to-dry weight ratio was measured as an indicator of pulmonary edema. Left lungs of a separate group of animals were explanted after reperfusion and patted dry. Lungs were weighed and then desiccated at 54°C under vacuum until a stable (dry) weight was obtained. Wet-to-dry weight ratios were calculated.

### Neutrophil immunohistochemistry

In separate groups of animals, left lungs were explanted after reperfusion, formalin-fixed via tracheal instillation, paraffin-embedded, and sectioned at 5-μm thickness. Following antigen retrieval in a microwave, neutrophils were identified by immunostaining using a rat anti-mouse neutrophil antibody (LY6B.2; Abcam). Alkaline phosphatase-conjugated anti-rat immunoglobulin G (Sigma-Aldrich) was used as a secondary antibody, and immune signals were detected using Fast-Red (Sigma-Aldrich). Purified normal rat immunoglobulin G (eBioscience, San Diego, CA) was used as a negative control. Sections were counterstained with hematoxylin. For each lung section, neutrophils were counted in five random fields at 20X magnification and averaged. These cell counts did not distinguish between cells in various compartments of the lung (e.g., airspace, interstitial, or marginated) but included all cells in peripheral (alveolar) lung tissue.

### Ca^2+^ imaging in Pas

Ca^2+^ imaging in small PAs was performed as previously described (*10*). Fourth-order, small (∼50 µm diameter) PAs (see Fig. 1B) were dissected from left lungs in cold HEPES-buffered physiological salt solution (HEPES-PSS; 10 mM HEPES, 134 mM NaCl, 6 mM KCl, 1 mM MgCl_2_ hexahydrate, 2 mM CaCl_2_ dihydrate, and 7 mM dextrose; pH adjusted to 7.4 using 1 M NaOH). Ca^2+^ imaging studies were performed in *en face* preparations of small PAs incubated with Fluo-4 AM (10 µM) and pluronic acid (0.04%) at 37°C for 10 minutes. For studies after sham or lung IR surgery, small PAs were immediately dissected out after the indicated reperfusion time and incubated *en face* with Fluo-4 AM. For acute hypoxia-reoxygenation (HR) exposure, small PAs were washed with PBS, after which serum free/glucose-free endothelial cell medium was added. Small PAs were placed in a humidified, sealed hypoxia chamber (Billups-Rothenberg, San Diego, CA), which was subsequently flushed with 95% nitrogen/5% CO_2_ for 20 minutes and placed in a 37°C incubator to initiate hypoxia. After 1 hour of hypoxia, the chamber was opened to terminate hypoxia. PAs were then washed with PBS; endothelial growth medium containing 10 μM Fluo-4 AM at 37°C was added, and PAs were reoxygenated by placing in a 37°C, humidified 5% CO_2_ incubator for 10 minutes. Ca^2+^ images were acquired at 30 frames per second using an Andor Revolution WD (with Borealis) spinning-disk confocal imaging system (Andor Technology, Belfast, UK) comprising an upright Nikon microscope with a 60X water-dipping objective (numerical aperture [NA], 1.0) and an electron-multiplying CCD camera (iXon888), as was done previously.(*19*) PAs were superfused with PSS (pH 7.4), and all experiments were performed at 37°C. Fluo-4 was excited using a 488-nm solid-state laser, and emitted fluorescence was captured using a 525/36-nm band-pass filter. PAs were treated with the sarco-endoplasmic reticulum (SR/ER) Ca^2+^-ATPase inhibitor cyclopiazonic acid (CPA; 20 µM) for 10 minutes at 37°C prior to imaging to eliminate intracellular Ca^2+^-release signals, respectively. We previously showed that CPA *per se* does not alter the activity of TRPV4 sparklets in endothelial cells. The effects of GSK219, ARC, and apyrase on pulmonary endothelial Ca^2+^ signals were studied 10 minutes after the addition of these drugs. Baseline Ca^2+^ signaling activity was recorded in the absence of CPA.

### Analysis of baseline Ca^2+^ signaling activity and TRPV4 sparklet activity in the pulmonary endothelium

Ca^2+^ images acquired using the procedure described above were analyzed with custom-designed SparkAn software (source code available at https://github.com/vesselman/SparkAn) (*24*). Fractional fluorescence (F/F_0_) traces were obtained by placing a 1.7 µm^2^ (5 × 5 pixels) region of interest (ROI) at the peak event amplitude. Representative F/F_0_ traces were filtered using a Gaussian filter and a cutoff corner frequency of 4 Hz. Overall baseline Ca^2+^ signals were automatically detected using a threshold of 1.25 F/F_0_. Changes in baseline (e.g., due to movement of the artery or a change in focus) were accounted for by obtaining a continuous baseline using a running-average approach from start to end, 19 points at a time, and then dividing the F/F_0_ trace by the continuous baseline. Each trace shown in figures indicates Ca^2+^ signaling activity at a single site.

TRPV4 sparklets were assessed as increases in fluorescence relative to baseline fluorescence, obtained by averaging 10 quiescent images prior to stimulation. TRPV4 Ca^2+^ sparklets were assessed based on previously established methods (*19, 46*). Average TRPV4 sparklet activity is defined as NP_O_, where N is the number of TRPV4 channels per site and P_O_ is the open-state probability of the channel. NP_O_ was calculated using the Single Channel Search module of Clampfit, quantal amplitudes derived from all-points histograms (0.3 ΔF/F_0_ for Fluo-4-loaded PAs) (*11*), and the following equation: NP_O_ = (*T_level1_* + *2T_level2_* + *3T_level3_* + *4T_level4_)/T_total_*, where *T* represents the dwell time at each quantal level and *T_total_* is the total recording duration. Average NP_O_ per site was obtained by averaging the NP_O_ for all sites in a field of view (110 μm x 110 μm). NP_O_ values per site from all fields in an artery were averaged to obtain NP_O_ per site for that artery. The total number of sites per cell corresponds to all sparklet sites in a field divided by the number of cells in that field. Sparklet sites per cell were averaged for all fields in an artery to obtain sparklet sites per cell for that artery.

### Exposure of PMVECs to acute HR and Ca^2+^ imaging in PMVECs

Primary human and murine pulmonary microvascular endothelial cells (PMVECs; Cell Biologics, Chicago, IL) were cultured on gelatin-coated flasks in endothelial cell medium (Cell Biologics). For acute HR exposure, PMVECs were washed with PBS, and serum-free/glucose-free endothelial cell medium was added. PMVECs were placed in a humidified, sealed hypoxia chamber (Billups-Rothenberg, San Diego, CA), whereupon the chamber was flushed with 95% nitrogen/5% CO_2_ for 20 minutes and placed in a 37°C incubator to initiate hypoxia. After 1 hour of hypoxia, the chamber was opened to terminate hypoxia. Cells were washed with PBS; complete endothelial growth medium containing 0.1 μM Calbryte 520 AM (AAT Bioquest, Pleasanton, CA) was added; and cultures were incubated at 37°C in a humidified 5% CO_2_ incubator (reoxygenation) for 10 minutes. Control, normoxic PMVECs underwent the same procedure except that the chamber was not flushed with nitrogen/CO_2_.

Ca^2+^ imaging in human and mouse PMVECs was performed using an Andor Dragonfly 505 spinning-disk confocal imaging system with Borealis illumination and an Integrated Laser Engine solid-state laser source (Andor Technology, Belfast, UK), coupled with a Leica DMi8 inverted microscope (Leica) and 63X objective (NA 1.47) (*21*). Images were acquired at 30 frames per second using Fusion 2.3 (Oxford Instruments, Belfast, UK) and an electron-multiplying CCD camera (iXon888). Calbryte 520 AM was excited at 488 nm, and emitted fluorescence was captured using a 525/50 bandpass emission filter. Image analysis was performed as described above for small PAs.

### Whole-cell patch-clamp electrophysiology

Patch-clamp electrodes were pulled with a Narishige PC-100 puller (Narishige International USA, Inc, Amityville, NY) and polished using a MicroForge MF-830 polisher (Narishige International, USA). The pipette resistance was (4-6 MΩ). Data were acquired using a Multiclamp 700B amplifier connected to a Digidata 1550B system and analyzed using Clampfit 11.1 software (Molecular Devices, San Jose, CA). TRPV4 channel currents were recorded from human and murine PMVECs. GSK101 (10 nM)-induced outward currents through TRPV4 channels were assessed using the whole-cell patch-clamp configuration in the presence of ruthenium red (RuR), included to prevent Ca^2+^ influx through TRPV4 channels and activation of potassium channels (*23, 46*). The intracellular solution consisted of 10 mM HEPES, 30 mM KCl, 10 mM NaCl, 110 mM K-aspartate, and 1 mM MgCl_2_ (adjusted to pH 7.2 with NaOH). HEPES-PSS was used as the bath solution. TRPV4 currents were measured using a voltage-clamp protocol in which voltage-ramp pulses (−100 mV to +100 mV) were applied over 200 ms from a holding potential of −50 mV. Currents were measured before and 5 minutes after treatment with GSK101 (10 nmol/L) followed by GSK219 (100 nmol/L), with GSK219-sensitive currents defined as TRPV4 channel-mediated currents.

### Quantitative polymerase chain reaction (qPCR)

Endothelial cell (EC) sheets were used for mRNA isolation, as described previously (*19*). Briefly, small PAs were enzymatically digested by incubating in dissociation solution (55 mM NaCl, 80 mM Na-glutamate, 6 mM KCl, 2 mM MgCl_2_, 0.1 mM CaCl_2_, 10 mM glucose, 10 mM HEPES, pH 7.3) containing 1 mg/mL neutral protease (Worthington, Freehold, NJ) for 30 minutes at 37°C. Following enzymatic digestion, EC sheets were gently squeezed out from small PAs with fine tips of 1-mL syringe needles. EC sheets were placed in a perfusion chamber and allowed to settle for 15 minutes at room temperature. EC sheets were collected with a micropipette (∼0.5–0.8 MΩ) by applying gentle suction. The purity of ECs (>95%) was confirmed in control experiments by immunostaining with FITC-tagged CD31 (*10*). Isolated ECs (∼2000) were transferred to TRI Reagent (Zymo Research, CA) and homogenized using a Standard Micro-homogenizer (PRO Scientific Inc., Oxford, CT). Total RNA was isolated using a Direct-zol RNA Miniprep Kit (Zymo Research, CA), treated with an on-column DNA removal protocol, and converted to cDNA using an iScript cDNA Synthesis Kit (Bio-Rad, Hercules, CA). qPCR was performed on a Bio-Rad CFX96 qPCR Detection System using a reaction mix containing Bio-Rad 2x SYBR Green Master Mix, 200 nM 5’ and 3’ primers, and 20 nM cDNA. qPCR primers for Panx1, TRPV4, and GAPDH (internal control) were obtained from GeneCoepia Inc. (Rockville, MD). P2Y2R qPCR primers were obtained from Bio-Rad (Hercules, CA). Results were analyzed using the ΔΔCt method.

### Luciferase assay for total ATP release

Small PAs were cut open and pinned down *en face* on a Sylgard block. PAs were then placed in black, opaque 96-well plates and incubated in HEPES-PSS for 10 minutes at 37°C, followed by incubation with the ectonucleotidase inhibitor ARL 67156 (300 μmol/L; Tocris Bioscience, Minneapolis, MN) for 30 minutes at 37°C. A 50-μL aliquot of each sample was transferred to another black, opaque 96-well plate, and ATP was measured using an ATP Bioluminescence HSII kit (Roche Applied Science, Penzberg, Germany) (*8*). Briefly, 50 μL of the luciferase reagent, luciferin, provided in the assay kit (Roche Applied Science), was added to each well and, following a 5-second orbital mix, luminescence was recorded using a luminometer (FluoStar Omega). ATP concentrations were calculated by reference to an ATP standard curve.

### Studies in freshly isolated small PAs from human lungs. Studies in freshly isolated small PAs from human lungs

Deidentified human lung wedge samples were obtained from normal-looking areas of explanted lung during lung transplantation surgeries, in accordance with policies of the University of Virginia Institutional Review Board (IRB #19044). Informed consent was obtained after the nature and possible consequences of the studies were explained. Multiple small PAs (∼50 μm) were dissected and used for Ca^2+^ imaging experiments, as described previously (*10*).

### Statistical analysis

Results are presented as means ± SEM. An n-value of 1 was defined as one small PA in imaging experiments (Ca^2+^ imaging), one cell in patch-clamp experiments, and one mouse for ATP measurements, qPCR experiments, lung function, lung edema, neutrophil infiltration, and inflammatory cytokine assays. Data were obtained from at least three mice in experiments performed in at least two independent batches. Individual data points are shown for each dataset. Graphical representations of all data were prepared using CorelDraw Graphics Suite X7 (Ottawa, ON, Canada), and all data were statistically analyzed using GraphPad Prism 8.3.0 (Sand Diego, CA). Power analyses used to determine group sizes and study power (>0.8) were performed using GLIMMPSE software (α = 0.05; >20% change). Using this method, we estimated at least five cells per group for patch-clamp experiments, five small PAs per group for imaging experiments, and five mice for qPCR, ATP, lung function, neutrophil, and cytokine measurements. Data were analyzed using two-tailed, paired or independent t-test for comparisons of data collected from two different treatments, and one- or two-way ANOVA to investigate statistical differences among more than two different treatments. A Tukey correction was applied for multiple comparisons with one-way ANOVA, and a Bonferroni correction was performed for multiple comparisons with two-way ANOVA. Statistical significance was defined as a p-value < 0.05.

## Supplemental Material

Figures S1-S5

## Acknowledgements

We thank Drs. Brant Isakson, Cheikh Seye, and Wolfgang Liedtke for floxed Panx1, P2Y2R, and TRPV4 mice, respectively, and Sree Katragadda for helping with lung images.

## Author Contributions

SKS and VEL conceptualized the study. MK, HND, AB, and SKS performed calcium imaging experiments and analysis. MK, HND, and AB performed HR experiments in PAs and PMVECs. HQT performed IRI surgeries and assessments of lung function, lung edema, neutrophil infiltration, and cytokine measurements. AK performed genotyping. ZD performed eATP measurements. MK, HQT, and SKS made figures. PWC provided human lung samples. MK and SKS wrote the manuscript. VEL and HQT edited the manuscript.

## Data and materials availability

All data needed to evaluate the conclusions in the paper are present in the paper or the Supplementary Materials.

