## Supplemental Material for "Purinergic P2Y2 Receptor-Induced Activation of Endothelial TRPV4 Channels Mediates Lung Ischemia-Reperfusion Injury"

### Endothelial Purinergic P2Y2 Receptor Signaling Mediates Lung Ischemia-Reperfusion Injury Through TRPV4 Channel Activation

Maniselvan Kuppusamy[1], Huy Q. Ta[2], Hannah N. Davenport[1], Abhishek Bazaz[1], Astha Kulshrestha[1], Zdravka Daneva[1], Yen-Lin Chen[1], Philip W. Carrott[2], Victor E. Laubach[2], Swapnil K. Sonkusare[1,3,*]

[1]Robert M. Berne Cardiovascular Research Center, University of Virginia, Charlottesville, VA 22908; [2]Department of Surgery, University of Virginia, Charlottesville, VA 22908; [3]Department of Molecular Physiology and Biological Physics, University of Virginia, Charlottesville, VA 22908

**Short Title:** Purinergic signaling in lung IR injury

*Corresponding Author:

Swapnil K. Sonkusare, Ph.D.

409 Lane Rd; room 6051A

Charlottesville, VA 22901

A

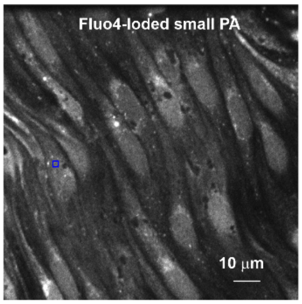


B

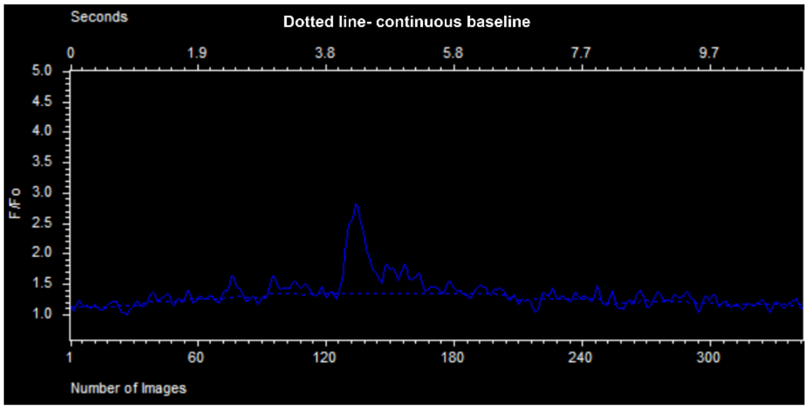


C

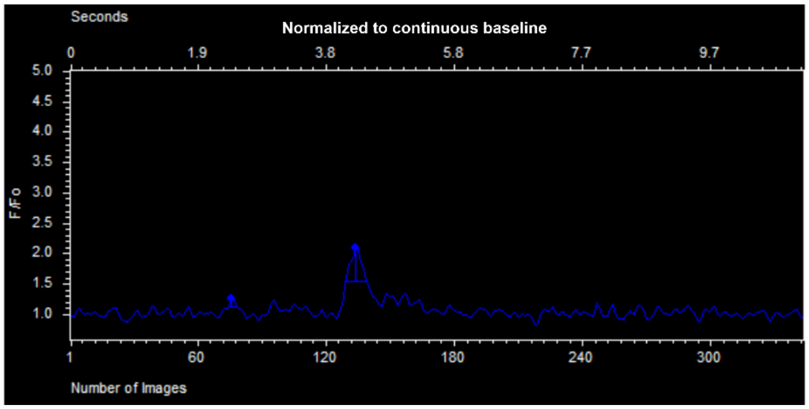


**Figure S1. Analysis of baseline $Ca^{2+}$ signaling activity in endothelial cells.** $Ca^{2+}$ images were analyzed using custom-designed SparkAn software. (A) Fractional fluorescence traces ($F/F_0$) were generated from regions of interest (ROI, blue square) placed at the peak event amplitudes. A continuous baseline was obtained for the duration of the recording (dotted line in B). Peaks were automatically detected after dividing the $F/F_0$ trace with the continuous baseline and using a threshold of 1.25 $F/F_0$ (C).

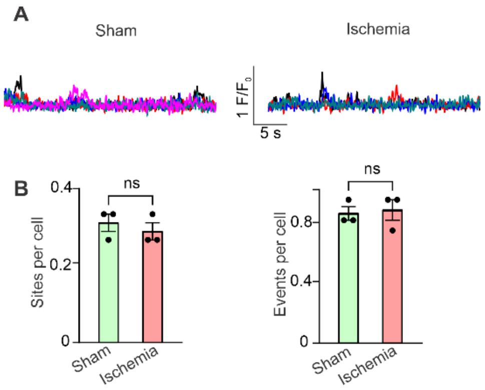


**Figure S2. Ischemia alone does not increase endothelial $Ca^{2+}$ signaling activity in small PAs.** (A) $F/F_0$ traces showing baseline $Ca^{2+}$ signaling activity from small PAs exposed to sham surgery or 1 hour of ischemia (no reperfusion). (B) Total endothelial $Ca^{2+}$ signaling activity represented as sites per cell and events per cell in small PAs (n = 3/group; ns, not significant; t-test).

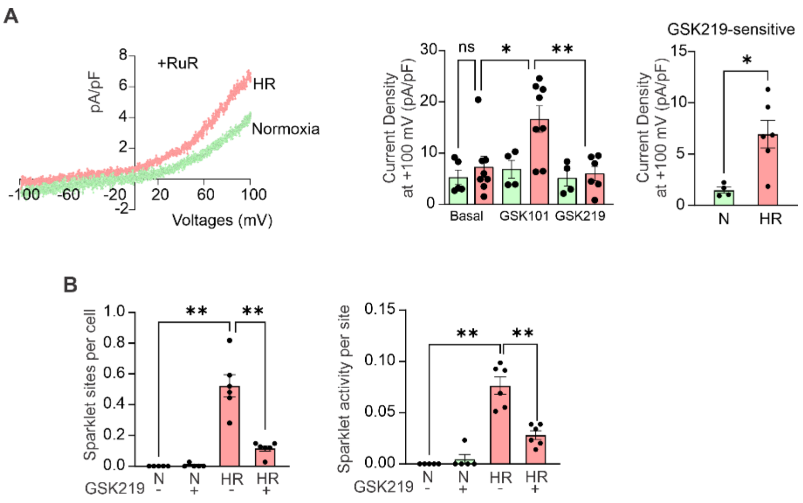


**Figure S3. Exposure of mouse PMVECs to acute HR induces endothelial TRPV4 channel activity.** Mouse PMVECs were exposed to 1 h hypoxia followed by 10 minutes reoxygenation (HR) or normoxia (N) for 70 minutes. (A) GSK2193874 (GSK219, TRPV4 inhibitor, 100 nM)-sensitive currents through TRPV4 channels in mouse PMVECs under normoxia and HR conditions recorded in the presence of RuR (1 μM, left) and dot plot showing outward currents at +100 mV (right). (B) TRPV4 sparklet activity (sites per cell, left; activity per site, right) after normoxia or HR exposure of mouse PMVECs in the absence and presence of GSK219 (n = 5-6/group; *P < 0.05; **P < 0.01; ns, not significant; two-way ANOVA).

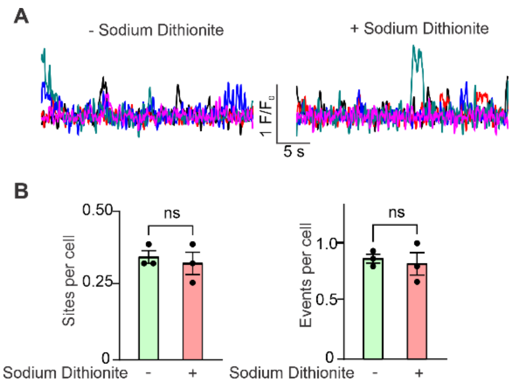


**Figure S4. Hypoxia alone does not increase endothelial $Ca^{2+}$ signaling activity in small PAs.** (A) $F/F_0$ traces showing baseline $Ca^{2+}$ signaling activity in small PAs exposed to normoxia or chemically-induced hypoxia (sodium dithionite, 1 mM). (B) Total baseline endothelial $Ca^{2+}$ signaling activity represented as sites per cell and events per cell (n = 3/group; ns, not significant; t-test).

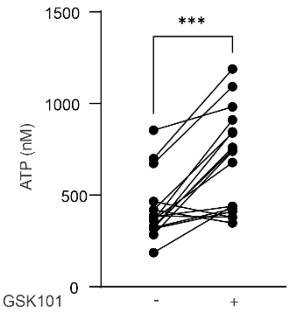


**Figure S5. Activation of TRPV4 channels with synthetic agonist GSK101 increases eATP release from small PAs** (A) eATP release (nM) from small PAs before and after addition of TRPV4 agonist GSK101 (10 nM) (n = 10-14/group; **P < 0.001; paired t-test).
